# Fundamental Structure and Modulation of Neuronal Excitability: Synaptic Control of Coding, Resonance, and Network Synchronization

**DOI:** 10.1101/022475

**Authors:** Christoph Kirst, Julian Ammer, Felix Felmy, Andreas Herz, Martin Stemmler

## Abstract

Neuronal encoding and collective network activity depend on the precise mechanism for generating action potentials. A dynamic switch in this mechanism could greatly expand the functional repertoire of neurons and circuits. Here we show that changes in neuronal biophysics control a complex, yet fundamental, sequence of dynamic transitions in neuronal excitability in which neurons switch from integrators to resonators near the spike threshold, from simple voltage dynamics to the bistable co-existence of action potentials and quiescence, and from continuous class-I to discontinuous class-II firing rate encoding. Using multiple bifurcation theory, we prove that this transition sequence is universal in conductance-based neurons. Using dynamic-clamp and pharmacology, we show experimentally that an increase in leak conductance or application of the inhibitory agonist GABA can dynamically induce these transitions in hippocampal and brainstem neurons. Our results imply that synaptic activity can flexibly control resonance, excitability and bistability of neurons. In simulated neuronal networks, we show that such synaptically induced transitions provide a mechanism for the dynamic gating of input signals and the targeted synchronization of sub-networks with a tunable number of neurons.

**Significance:** Neuronal function depends on the mechanism by which neurons transform synaptic input into action potentials (APs). It is unclear how neurons might control the AP mechanism to systematically modulate their responses to input signals or their collective behavior. Here we identify a complex, but model-independent, universal sequence of transitions in the dynamics of AP generation. Using patch-clamp recordings, we show that synaptic receptor activation can flexibly change the AP dynamics, confirming our theoretical predictions: non-resonant neurons develop a sub-threshold resonance, become bistable, and develop an abrupt jump in onset AP frequency. Our results explain how synapses or neuro-modulators could control neuronal excitability, influence information processing, and processing during collective network dynamics.

## Introduction

Membrane potential dynamics and action potential (AP) generation are fundamental to neuronal coding, governing signal processing and network behavior throughout the nervous system. Systematic modulation of the dynamics at the single neuron level has the potential to flexibly control responses to stimuli as well as collective activity in networks. To what degree the voltage dynamics and AP generation can be actively changed to control these properties in single cells is an open question, however.

A marker for the type of AP dynamics is the transformation of injected current (*I*) via an output firing rate (*f*) as measured by a neuron’s *f* -*I*-curve [1, 2, 3]. Whether this curve rises continuously from zero (class-I neuronal excitability) or jumps discontinuously to a non-zero frequency (class-II) strongly influences the neuron’s selectivity to stimulus features [2, 3, 4], the precision of spike timing [5, 6, 7, 8], information processing and storage [9, 10] as well as the collective dynamics on the network level [11, 12, 13]. Class-II excitability often, but not exclusively, arises through the amplification of damped membrane voltage oscillations. In this case, the neuron resonates to oscillatory input even before reaching the threshold for APs. Such membrane potential resonances facilitate participation in network oscillations [14, 15, 16, 17] and shape signal transduction [18, 19].

Both neuronal excitability and resonance properties are governed by many factors, including *I*_h_, *I*_M_, or adaptation currents [20, 21, 22], neuromodulators [21, 23], *in vivo* vs. *in vitro* conditions [24], or even a neuron’s input resistance [2, 20, 24]. It is not clear, though, whether there is a one-to-one correspondence between neuronal excitability and resonance, what higher-order bifurcations are triggering transitions in AP generation, and whether synaptic neurotransmitter release can actively induce these transitions.

## Results

### Leak-Induced Transitions in AP Dynamics

To study how AP generation can be modulated, we consider conductance-based neuron models of the form

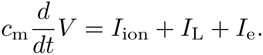

Here *V* is the membrane potential, *c*_m_ the neuron’s capacitance, *I*_ion_ represents trans-membrane currents of active ion channels, *I*_e_ is an externally applied current. We first focus on how the leak conductance *g*_L_, a property generic to all neurons, affects neuronal excitability. This conductance gives rise to a leak current

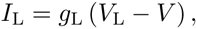

with reversal potential *V*_L_.

Increasing the leak conductance switches class-I model neurons to class-II (Fig. 1A), as the initially continuous *f* -*I*-curve becomes step-like. Class-I spiking originates from a saddle fixed-point (SI Appendix, Lemma S5) with stable and unstable directions of the dynamical flow (Fig. 1A, first sketch). Along the unstable direction, large voltage excursions emerge that represent APs. As *I*_e_ is increased, the saddle merges with the resting state, and the loop in the dynamics closes in on itself, yielding a stable limit cycle of periodic AP generation (Fig. 1A, second sketch) – known as a saddle-node on limit cycle (SNLC) bifurcation [2, 12]. Near the saddle, the dynamics slows down, permitting arbitrarily long inter-spike intervals. In contrast, class-II spiking often reflects the destabilization of small-amplitude oscillations as *I*_e_ increases.

**Fig. 1:**
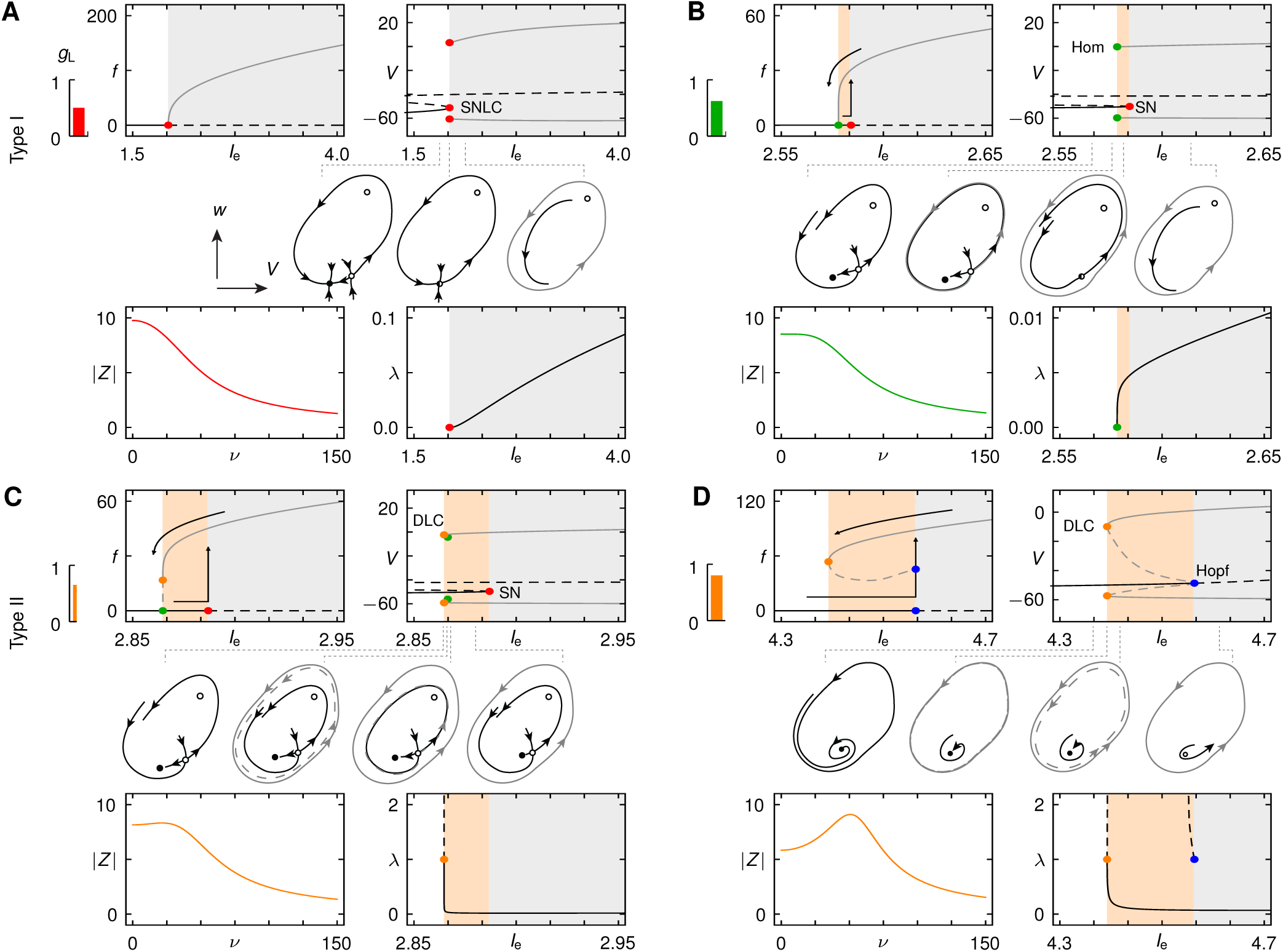
Leak-induced transitions in neuronal excitability and resonance. Each panel shows the *f* -*I*-curve (top left), the voltage *V* against the input current *I*_e_ (top right) and sketches of the AP dynamics in a two-dimensional phase-plane of *V* and an effective activation variable *w* (middle), the impedance |*Z*| at 95% of the current at Ap threshold (bottom left) and the largest non-trivial Floquet multiplier (FM) *λ* (bottom right) for different values of the leak conductance *g*_L_ in the Wang-Buzsáki neuron model. Black lines –(-) indicate (un)stable fixed points, gray shading 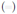 stable periodic spiking, orange shading 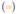 bistability in which a steady state and spiking coexist. Gray lines (–) denote the maximum and minimum voltages of the stable AP cycle, dashing (-) indicates unstable orbits. In the sketches, closed (open) dots show stable (unstable) fixed points, black lines stable and unstable manifolds, gray lines periodic orbits. (*A*) For small *g*_L_, a saddle-node on limit cycle bifurcation (SNLC, 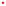) generates a continuous *f*-*I*-curve, the hallmark of class-I_*a*_ excitability. *Z* decays steadily, so there is no resonance and small FM reflects fast attraction towards the limit cycle. (*B*) For intermediate *g*_L_, the neuron shows a mixture of class-I and II excitability (I_b_). A stable limit cycle arises through a homoclinic bifurcation (Hom, 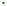); the resting state is destabilized for higher *I*_e_ in a saddle-node bifurcation (SN, 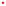). The resulting hysteresis in firing rates shows non-zero spike-frequency onset but a gradual decay to zero at offset. There is no resonance; the FM is zero at the bifurcation point. (*C*) For even larger *g*_L_, class-II_*a*_ dynamics appear. A double limit cycle bifurcation (DLC, 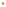) generates AP cycles with finite period, resulting in a discontinuous *f* − *I*-curve. The unstable cycle vanishes in a homoclinic bifurcation (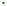), destabilization of the resting state is through a saddle-node, as before. saddle-node, as before. There is a small resonance close to threshold. At the bifurcation, the FM is unity, so attraction to the AP limit cycle is slow. (*D*) For large *g*_L_, the fixed point is destabilized via a Hopf bifurcation (Hopf,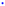). The resonance near threshold becomes more pronounced.

In class-I neurons, the dynamics converge quickly towards the limit cycle. The convergence is measured by a non-trivial Floquet-multiplier, which is zero when the dynamics converges towards the limit cycle within one period and unity when there is no attraction. In class-I neurons, this multiplier is close to zero (Fig. 1A, bottom right).

At the saddle-node point the linear membrane-potential response amplitude *Z* (*ν*) to sinusoidal current stimuli with frequency *ν* becomes proportional to 1*/ιν* at low frequencies (Fig. 1A, bottom left). In frequency space, multiplication by 1*/ιν* is equivalent to integration, which implies that the neuron integrates current inputs until the AP threshold is reached.

As *g*_L_ increases, the limit cycle splits off directly from the saddle (Fig. 1B) in a homoclinic bifurcation [3], while the stable fixed point remains. As a consequence, a region of bistability and hysteresis in the *f* -*I*-curve emerges. We term the mixture between class-I for AP-offset and class-II for AP-onset class-I_*b*_, to discriminate it from class-I_*a*_ that is continuous in both directions.

For higher *g*_L_, a stable and unstable periodic orbit arise through a double limit cycle (DLC) bifurcation, while the saddle-node bifurcation continues to mark the destabilization of the resting state (Fig. 1C). Both AP on-and offset are discontinuous (class-II_*a*_), the frequency response |*Z* (*ν*)| shows a resonance, and a non-zero Floquet multiplier implies a slower decay of the spike amplitude. Increasing *g*_L_ further, the saddle-node gives way to a sub-critical Hopf bifurcation (Fig. 1D, top) to destabilize the resting state. Damped oscillations become unstable, resulting in a strong resonance (Fig. 1D, bottom left).

Finally, the DLC and sub-critical Hopf bifurcations coalesce and turn into a super-critical Hopf bifurcation, while the region of bistability vanishes. The *f* -*I*-curve becomes class-II without hysteresis (class-II_*b*_).

### Fundamental Structure of Neuronal Excitability

The full structure of the transitions becomes visible in the two-dimensional bifurcation diagram in (*I*_e_,*g*_L_)-parameter space (Fig. 2A): two large regions correspond to the two main excitability classes observed in Fig. 1A,D. Enlarging the transition region (Fig. 2A) reveals the bifurcations that mark the individual changes (cf. SI Appendix, Fig. S1, S2 for phase-plane dynamics): a saddle-node-loop (SNL) bifurcation separates class-I_*a*_ excitability (SNLC, Fig. 1A) from the mixture class-I_*b*_ that supports dynamical bistability (homoclinic, Fig. 1B). The switch from class-I_*b*_ to II_*a*_ (DLC, Fig. 1C,D) occurs at a neutral saddle loop (NSL) point. A degenerate Hopf (DH) point controls the final change from class-II_*a*_ to II_*b*_, during which the bistability disappears. Destabilization of the resting state occurs first through a saddle-node (Fig. 1A-C) and then through a Hopf bifurcation (Fig. 1D) above the Bogdanov-Takens (BT) point.

**Fig. 2:**
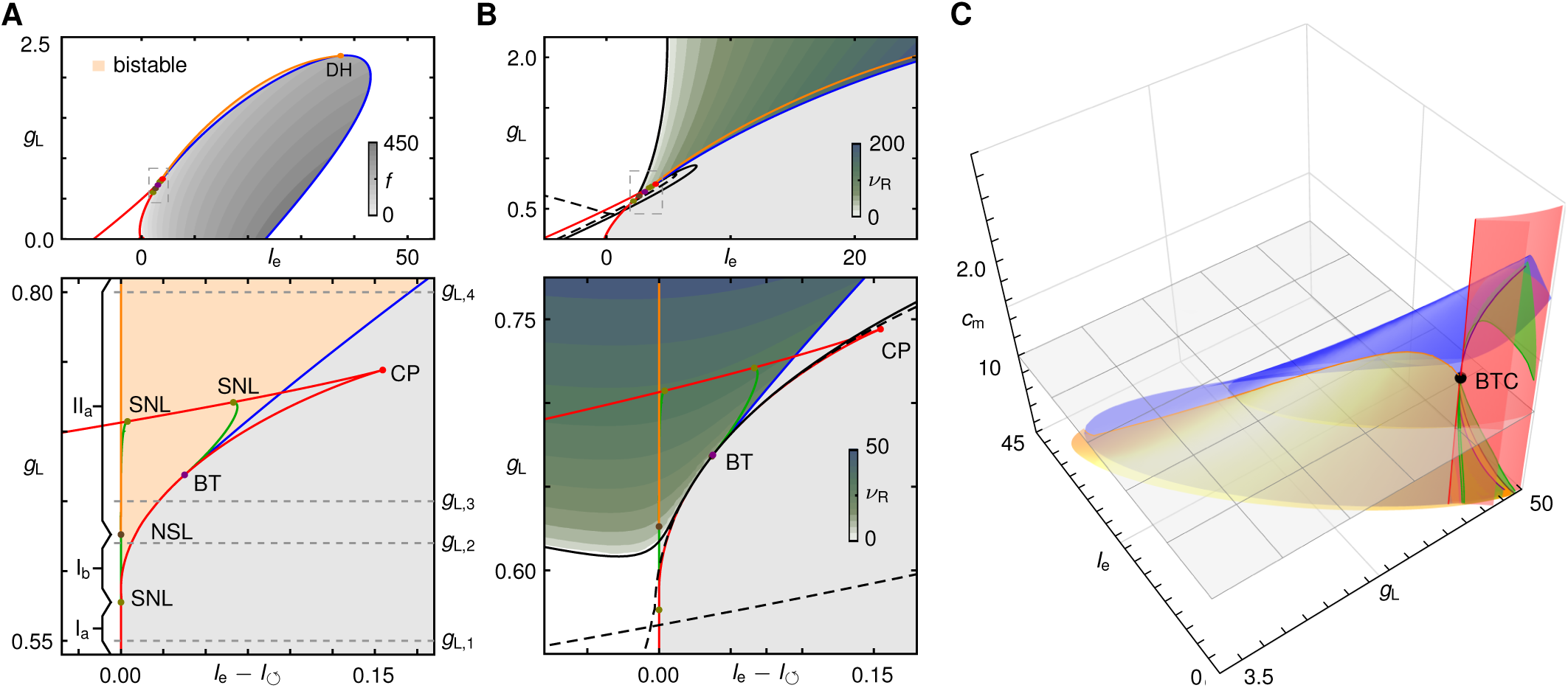
Fundamental structure of neuronal excitability transitions. Bifurcation diagram for the neuron model in Fig. 1 in the input current *I*_e_ and leak *g*_L_ parameter plane. Co-dimension one bifurcations: saddle-node 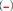, homoclinic 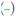, double limit cycle (DC,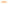), and Hopf 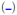 bifurcations. Gray level 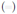 represents the frequency of periodic APs, orange shading 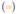 bi-stability between resting an spiking. Co-dimension two bifurcations: Bogdanov-Takens (BT, 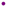), cusp (CP,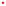), degenerate Hopf (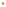), saddle-node loop (SNL, 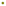) and neutral saddle loop (NSL,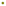). Top, full diagram. Bottom, fine structure of the transition with input current *I*_e_ is shifted by the onset current for stable spiking 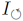. Dashed horizontal lines refer to *g*_L_ values used in Fig. 1. Class-I_a_ excitability due to a SNIC 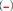, changes via the SNL-point (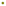) to class-I_b_ governed by a homoclinic bifurcation 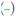; a NSL-point 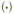 marks the transition to class-II_a_ (DC,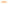). A degenerate Hopf bifurcation separates class-II_a_ from class-II_b_ behavior (visible only in (A)). A region of bistability 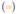 emerges at a SNL (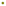) and vanishes at the degenerate Hopf point. At the BT-point (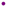), destabilization of the resting state changes from saddle-node 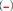 to Hopf 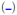. (*B*) Leak-induced resonance near AP-threshold. Colors and panels as in (A), green shading encodes resonance frequencies *ν*_R_ near AP-threshold. At the black dashed line (--) the real eigenvalues of the linearized dynamics become imaginary, indicating onset of oscillatory components in the response. Black solid line (–) shows the transitions from zero to non-zero resonance frequency *ν*_R_. The transitions to resonance (–,--) pass tangentially through the BT-point (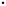). For *g*_L_ below the BT-point and increasing *I*_e_, the resonance frequency *ν*_R_ first increases, then dips just before the AP threshold is reached. (*C*) Neuronal excitability transitions are organized by a Bogdanov-Takens-cusp bifurcation (BTC, 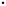) in (*I*_e_,*g*_L_,*c*_m_)-parameter space. Surfaces are co-dimension one (colors as lines in (A)) and lines are co-dimension two bifurcations (colors as points in (A)). The surface at *c*_m_ = 1 shows the plane of the diagrams in (A). At the BTC-point, the complex bifurcation diagram collapses into a single point that organizes the transition structure.

In addition to the transitions in neuronal excitability and towards bistability (Fig. 2A) membrane-potential resonances are observed in a wide swath of (*I*_e_,*g*_L_)-parameter space (Fig. 2B). As *g*_L_ increases, so does the resonance frequency *ν*_0_. Intriguingly, our mathematical results show that resonance does not imply that damped membrane potential oscillations at threshold are amplified to yield APs, as predicted by a Hopf bifurcation. Indeed, neurons can display both peri-threshold resonance and class-I AP dynamics: Below the BT point in the (*I*_e_,*g*_L_)-plane, the resonance frequency first increases with *I*_e_, but then rapidly decays back to zero before the AP current threshold inherent in the saddle-node bifurcation of the resting state (Fig. 2B, SI Appendix, Fig. S3). Interestingly, these resonances are solely a property of the AP generator and no specialized currents such as *I*_h_, *I*_M_, or *I*_Nap_ are necessary to induce them. While AP-independent sub-threshold oscillations are attenuated by leak [14, 29], here increasing the leak conductance is a prerequisite for resonances to appear.

The bifurcation diagrams in Fig. 2A,B are prevalent in many neuron models (Methods, SI Appendix, Figs. S4-S6, Table S1). Indeed, there is a generic mechanism underlying the sequence of transitions: Including the membrane capacitance *c*_m_ as a third generic parameter in the bifurcation analysis, we obtain a three-dimensional (*I*_e_,*g*_L_,*c*_m_)-parameter diagram (Fig. 2C). Remarkably, a combination of parameter values exists for which the transition structure collapses into a single degenerate point: a Bogdanov-Takens-cusp (BTC) bifurcation [25]. Using a combination of a center-manifold and normal form reduction [26] we prove that every class-I conductance-based neuron model has such a BTC point (Methods, Theorem) and give conditions for its particular subtype to be either focus or elliptic (Methods, Proposition). The proof relies on the fact that the bifurcation parameters *i*_e_, *g*_L_, and *c*_m_, are general parameters that appear in all conductance-based neuron models, and that in such models the dynamics of the gating variables are solely coupled to the membrane potential. According to multiple bifurcation theory [25, 27, 28], a focal or elliptic BTC point unfolds upon changing the parameters (*I*_e_,*g*_L_,*c*_m_) to yield the bifurcation structure observed in Fig. 2C. Taking a two-dimensional section of the diagram below the BTC point yields the leak-induced transition structure in the (*I*_e_,*g*_L_)-plane observed in Figs. 2A,B, SI Appendix, Figs. S1, S4-S6. The BTC point provides the link between the transitions in excitability and bistability (Fig. 2A) and the appearance of resonances (Fig. 2B). In addition, the normal form for the unfolding of the BTC point (Methods, Eq. (0.3)) provides as a simple generic neuron model that captures all aspects of the transition sequence.

While we prove that any class-I model has a BTC point, the converse is not necessarily true: for some class-II neuron models, the transition to class-I behavior would occur at negative leak conductances, which could cause the dynamics to become unbounded and are bio-physically implausible. (SI Appendix, Section 2).

The BTC transition structure persists in neuron models with firing rate adaptation (SI Appendix, Fig. S6). Adaptation decreases the minimal leak conductance required to induce the transitions, the region of bistability becomes broader, while spike and resonance frequencies are weakly affected.

The leak conductance is not the only parameter that induces a sequence of transitions typical for a BTC unfolding. Decreasing the maximal conductance of the fast depolarization currents, lowering the midpoint of the delayed rectifier current’s activation, or increasing the maximal conductance of the delayed rectifier all give rise to transition structures that are two-dimensional sections of the full three-dimensional BTC diagram (SI Appendix, Figs. S1, S2, S5). As a consequence, the reduced equations (0.3), generically capture all aspects of neuronal excitability transitions and may hence serve as a simple model to study effects of those modulations in excitability on collective network dynamics (also cf. below).

In summary, the BTC point and its unfolded bifurcation diagram are present in all class-I conductance based neuron models. The unfolding of the BTC point in turn predicts the precise sequence of dynamical transitions in response to any change that affects AP generation.

### Leak-Induced Neuronal Excitability Transitions in Vitro

Three main predictions follow from our theoretical results (Fig. 3): First, a region of bistability should emerge as the leak conductance is increased, so that noise fluctuations would drive the neuron to alternate between periodic spiking and sub-threshold oscillations (Fig. 3A,C). Second, the neuron should switch from peri-threshold integration to resonance (Fig. 3B,E,F). Third, the excitability should change from class-I to II (Fig. 3A). During this process, the maximal Floquet multiplier, measured at spike threshold, gradually changes from close to zero to close to unity (Fig. 1). As a consequence, class-I APs should maintain nearly constant amplitudes, whereas, for class-II, the AP trajectory decays slowly towards the stable limit cycle [3, 30] (Fig. 3C,D).

**Fig. 3:**
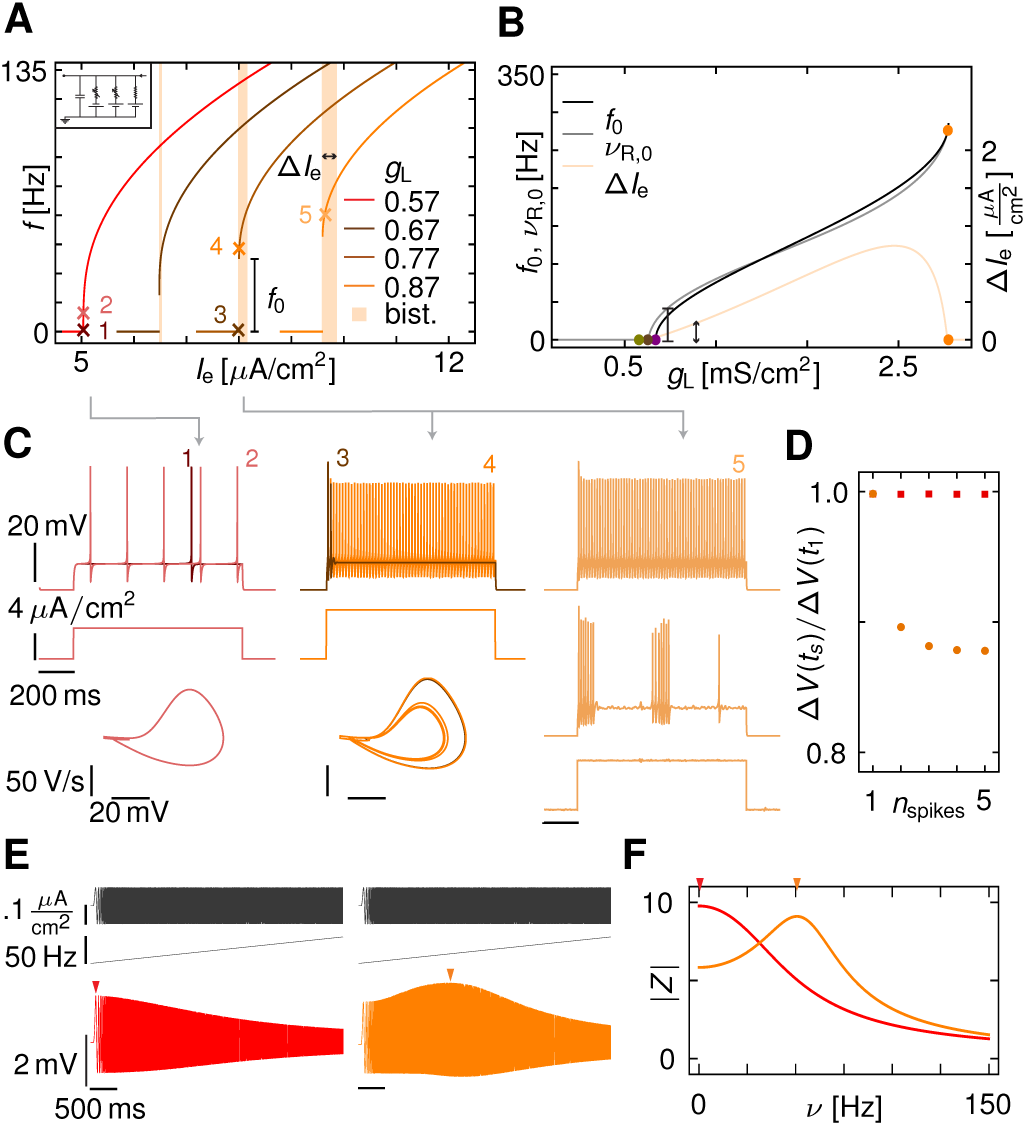
Leak-induced excitability transitions in firing rate encoding, resonance and AP characteristics. (A) The *f* -*I*-curves for increasing leak conductances in the model of Fig. 1 become discontinuous as the AP frequency jumps from 0 to *f*_0_, and regions of bistability 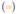 between quiescence and spiking emerge. (*B*) AP frequency *f*_0_ at threshold, resonance frequency *ν*_R_ (at 95% threshold current) and width Δ*I*_e_ of the bistable region as a function of *g*_L_. Scale bars as in (*A*). Colored dots indicate co-dimension two bifurcations that organize the individual sub-transitions as in Fig. 2. (*C*) Voltage traces (top) and phase portraits 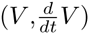 plane, bottom) of the AP dynamics in response to step currents (middle) for class-I (red) and II (orange) neuronal excitability (indicated by crosses in (A)). The amplitude dependence for class-II AP generation is clearly visible in the phase portrait. Parameters for the traces on the right are from the bistable region (cf. light orange crosses in (A)) for a deterministic step current (top) and a step current with added noise (middle, bottom) that induces switching between spiking and oscillatory fluctuations around the stable fixed point. (*D*) AP amplitudes for the traces in (C), which decay for class-II (orange), but remain stable for class-I (red). Floquet multipliers are measured to be 0.68 and 0.32, respectively. (*E*) Voltage responses (bottom) to ZAP stimuli (top) for class-I (left) and class-II excitability show a shift of the maximal response amplitude from 0 Hz (red arrow) to a non-zero frequency (orange arrow). (*F*) Impedance |*Z*| obtained from the responses in (E) change from input integration (red) to resonance (orange) with non-zero resonance frequency *ν*_R_ (orange arrow) as in (B).

We used dynamic patch-clamp recordings to test these predictions by artificially adding an external leak conductance *g*_L,e_ to the intrinsic leak *g*_L,0_ of the neurons (cf. also SI Appendix, Fig. S7A,B). The firing response to step-currents of a neuron in a slice from the dorsal nucleus of the lateral lemniscus (DNLL) in a Mongolian gerbil is shown in Fig. 4A for different externally applied leak conductances. For *g*_L,e_ = 0 nS the neuron exhibited slow firing close to AP threshold, indicating class-I behavior, supported by the near constancy of the AP amplitudes (Fig. 4D,E). An increase in leak conductance to *g*_L,e_ = 9, 20, and 30 nS induced a growing discontinuity in the onset frequency (Fig. 4B) and decaying AP amplitudes (Fig. 4D,E), indicating a switch to class-II excitability near *g*_L,e_ = 9 nS. At *g*_L,e_ = 30 nS, a region of bistability emerged (Fig. 4A,C, Methods).

**Fig. 4:**
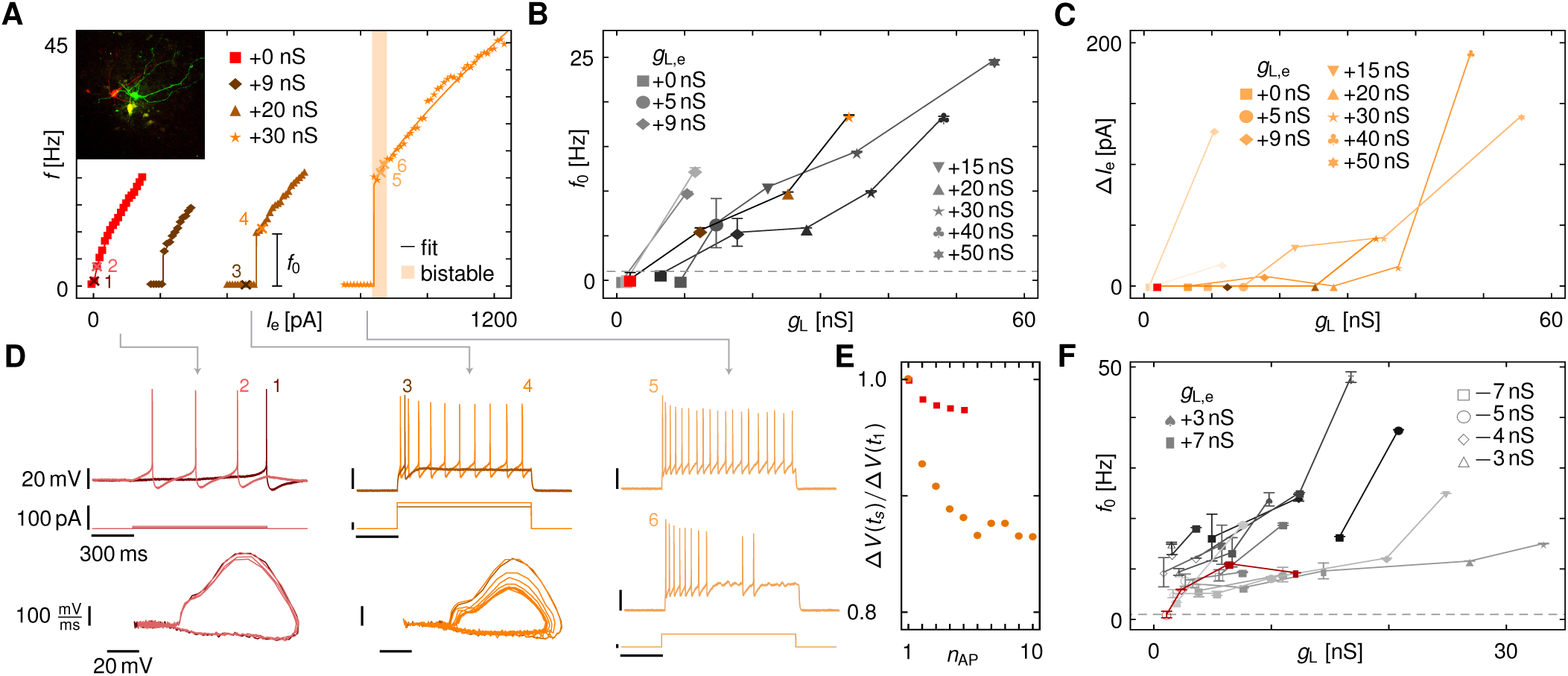
Leak-induced excitability and resonance transitions in the AP dynamics of DNLL neurons in vitro. (*A*) *f* -*I*-curves of a class-I DNLL neuron for different values of an added, artificial leak conductance that shifted the threshold for APs, switched the neuron from class-I to II and induced bistable dynamics 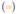. See Fig. 3 for model predictions. Crosses indicate the values for the traces shown in (D). Inset are stained DNLL neurons from recordings. (*B*) Increase in onset firing frequency *f*_0_ with increasing *g*_L,e_ across different class-I neurons (*n* = 5). The data from (A) is highlighted in color. (*C)* Width Δ*I*_e_ of the region of bistability for the cells in (B) systematically increases with leak. (*D*) Voltage traces (top) and phase portraits (bottom) show the dynamics in response to step-currents (middle) for *g*_L,e_ = 0 (left, red) and 20 nS (middle, orange). Right traces show two responses (top, middle) at *g*_L,e_ = 30 nS to two similar step current injections (bottom) in the region of bistability. Periodic firing (top) or switching dynamics between spiking and peri-threshold oscillations (middle) are observed. The traces are marked as crosses in (A). (*E*) AP voltage amplitudes vs. AP number *n*_AP_ within a response to a step-current for *g*_L,e_ = 0 (red) and 20 nS (orange) at AP threshold. Estimated Floquet multipliers are 0.37 and 0.69. (*F*) Onset firing rate *f*_0_ in intrinsically class-II DNLL neurons (*n* = 13). Increase of *g*_L,e_ increased firing frequency as in (B), subtraction of leak switched a single neuron to class-I (dark red); the dynamics of the remaining neurons became unstable upon large *g*_L,e_ subtraction.

We found similar transitions in all intrinsic class-I neurons (onset frequency < 1 Hz, stable AP amplitudes) measured in the DNLL (*n* = 5). Upon adding an artificial leak conductance, all these neurons switched to class-II excitability (Fig. 4B, S7D) and bistability was induced (Fig. 4C, S7E). The monotonic increase in onset AP frequency *f*_0_ and the width Δ*I*_e_ of the bistable region as a function of the measured leak *g*_L_ parallels those predicted theoretically (Fig. 3B). The switch in excitability is corroborated by a rapid decay of the latency to spike onset as the leak conductance is increased (SI Appendix, Fig. S7F,G). Application of very high leak conductances induced silencing of AP generation via a Hopf bifurcation (SI Appendix, Fig. S7C) in further agreement with a BTC transition scenario.

Most of the measured DNLL neurons showed intrinsic class-II excitability (*n* = 13*/*18) with systematically increasing onset frequency *f*_0_ as *g*_L_ increases (Fig. 4E). Subtraction of leak led to smaller onset frequencies, but a switch to pure class-I behavior was observed in only one neuron; in other neurons such a switch did not occur before the subtraction of leak caused the recording to become unstable. This finding is consistent with our theory not guaranteeing the reverse switch from class-II to I excitability.

To test the predicted generality of our results, we performed the same experiments in hippocampal CA3 pyramidal neurons (Fig. 5). The *f* -*I*-curves and the phase-plane dynamics showed a shift in neuronal excitability and a region of bistability, as predicted by theory (Fig. 5A,D,E). All intrinsically class-I neurons (*n* = 12) exhibited this behavior (Fig. 5B). The range of applied currents Δ*I*_e_ for which bistable spiking and quiescence were observed was larger than in the DNLL neurons, also increased with leak (Fig. 5C) while the latency to the first spike rapidly decreased (Fig. 5F). For intrinsically class-II neurons (*n* = 6) subtracting leak decreased the AP frequency and the width of the bistable region (SI Appendix, Fig. S7H), but did not change the dynamics to class-I. Intrinsic leak also correlated with intrinsic onset frequency in both DNLL (*n* = 23, *r* = 0.76) and CA3 neurons (*n* = 18,*r* = 0.77, SI Appendix, Fig. S7J,K).

**Fig. 5:**
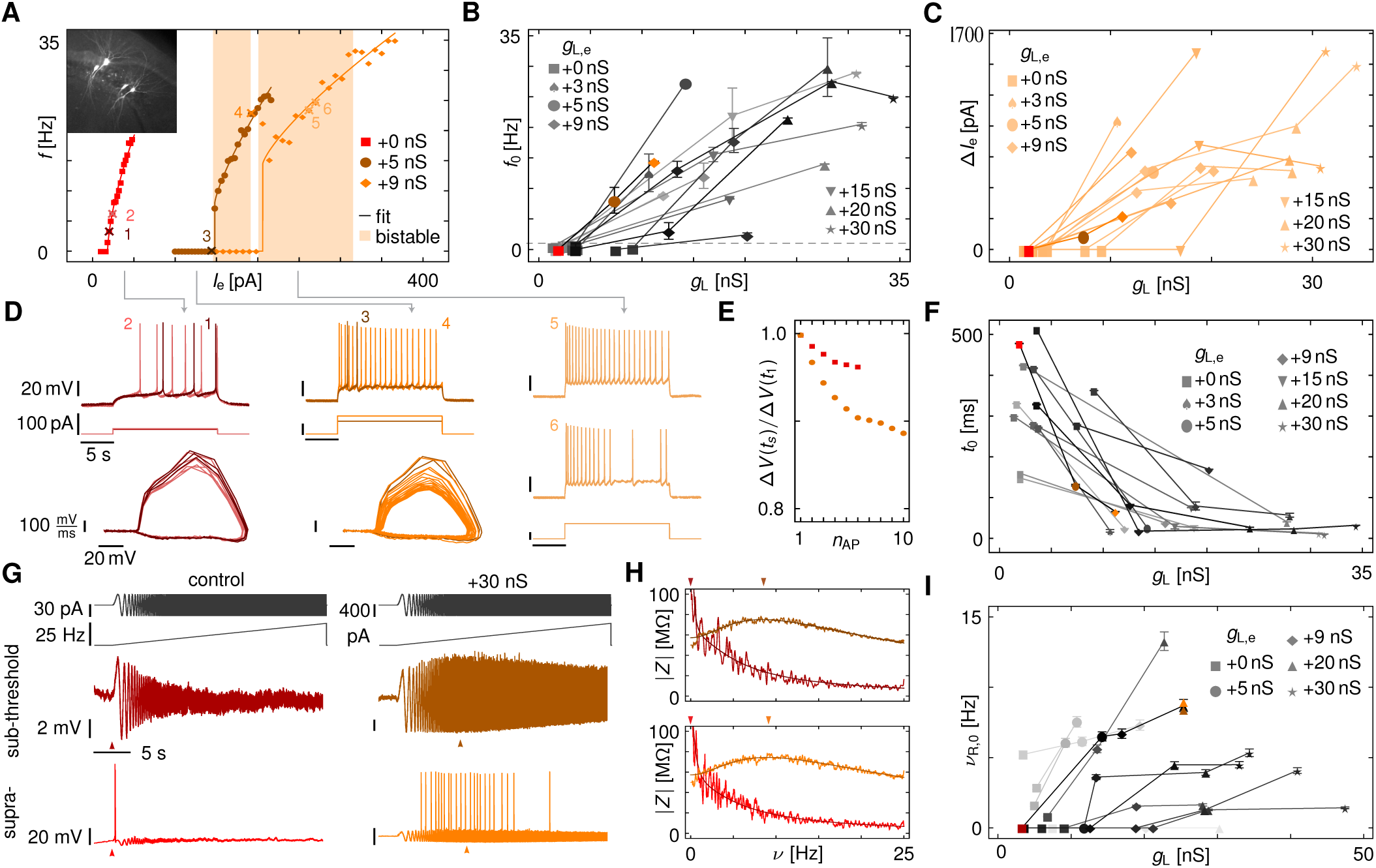
Leak-induced excitability and resonance transitions in the AP dynamics of CA3 neurons in vitro. (*A*) *f* -*I*-curves of a class-I CA3 neuron for increasing values of an added, artificial leak conductance show increase in onset firing frequency *f*_0_ and induction of bistable dynamics 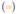. (*B*) Increase in onset firing frequency with increasing *g*_L,e_ across different class-I neurons (*n* = 12). The data from (*A*) is highlighted in color. (*C)* Width Δ*I*_e_ of the region of bistability for the cells in (B) systematically increases with leak as predicted by theory (cf. Fig. 3B). (*D*) Voltage traces (top) and phase portraits (bottom) show the dynamics in response to step-currents (middle) for *g*_L,e_ = 0 (left, red) and 5 nS (middle, orange). Right traces show two responses (top, middle) at *g*_L,e_ = 9 nS to two similar step current injections (bottom) in the region of bistability. The traces are marked as crosses in (A). Periodic firing (top) or switching dynamics between spiking and peri-threshold oscillations (middle) are observed. (*E*) AP voltage amplitudes vs. AP number *n*_AP_ within a response to a step-current for *g*_L,e_ = 0 (red) and 9 nS (orange). (*F*) Latency *t*_0_ from step stimulus onset to first AP in the class-I CA3 neurons in (B). Increased leak conductance strongly reduces the time to the first spike.(*G*) Voltage responses of a CA3 pyramidal cell to a ZAP current with linearly increasing frequency from 0 to 25 Hz (top) in 30 s. Responses just below (middle) and above spiking threshold (bottom) for *g*_L,e_ = 0 (left, red) and 30 nS (right, orange). (*H*) Impedance |*Z* (*ν*) | (line) for the traces in (G) with fit to an LRC circuit (darker line). Arrows indicate resonance frequencies *ν*_R_ obtained from the fit. (*I*) Average *ν*_R,0_ of sub-and supra threshold resonance frequencies *ν*_R_ as a function of *g*_L,e_ for all cells (*n* = 10), six of which did not intrinsically resonate. Cell in (G,H) highlighted in color. The resonance increases systematically with increasing leak, as predicted by theory (Fig. 2B, 3B,E,F).

To detect a resonance frequency close to AP threshold, we measured the neurons’ impedance from their response to injection of ZAP currents [31], which are oscillations whose frequency increases linearly with time. For an intrinsically class-I neuron, the voltage responses below threshold decreased as the frequency in the ZAP increased (Fig. 5G-I). Above threshold, a single AP was generated on the first peak in the stimulus. In both cases, the impedance curve decayed monotonically and had no detectable corner frequency, as expected for an integrator (*ν*_R_ = 0 Hz, Fig. 5G,H).

An increase to *g*_L,e_ = 30 nS changed the impedance profile and a resonance developed (Fig. 5G,H) below (*ν*_R_ = 8.6 Hz) and above threshold (*ν*_R_ = 9.1 Hz). In all non-resonant (*n* = 6) as well as intrinsically resonant neurons (*n* = 4), adding leak conductance increased the resonance frequency (Fig. 5I). The strength of the resonance (Q-factor) ranged between 1.1 and 3.2 (SI Appendix, Fig. S8A). The same behavior was observed in DNLL neurons (SI Appendix, Fig. S8E). We further observed a non-monotonic dependence of the resonance frequency as a function of the holding potential before AP threshold was reached (SI Appendix, Fig. S8C,D), as predicted by theory (SI Appendix, Fig. S3).

Dynamic-clamp recordings of both DNLL and CA3 neurons thus confirmed the theoretical predictions from the Bogdanov-Takens-cusp theory: leak induces a transition from class-I to class-II excitability, generates regions of bistable dynamics, and switches neurons from integration to resonance.

### GABA-Induced Excitability Transitions in CA3

We hypothesized that synaptic inhibition can increase the effective leak conductance and so permit synaptic control of neuronal excitability and resonance. To test this, we puffed 500 *μ*M GABA locally onto CA3 cells. This indeed induced a switch in neuronal excitability (*n* = 7, Figs. 6A,B) and towards bistability (Fig. 6C). Furthermore, activation of GABA also induced a switch from peri-threshold integration to resonance (Fig. 6D-F, SI Appendix, Fig. S9). GABA led to an increase in leak conductance of *g*_L,GABA_ = 5 *-* 10 nS as estimated from the cell’s response to small negative current steps. The changes in onset AP frequency, resonance frequency and quality factor (Fig. 6, SI Appendix Fig. 9) matched those found when artificially adding a similar amount of leak via dynamic clamp (Fig. 5, SI Appendix, S7H and S8B). These results suggest that activation of shunting synapses alone can dynamically move the neuron between different regimes in the transition diagram and thereby control the neurons’ resonance, bistability and excitability properties.

**Fig. 6:**
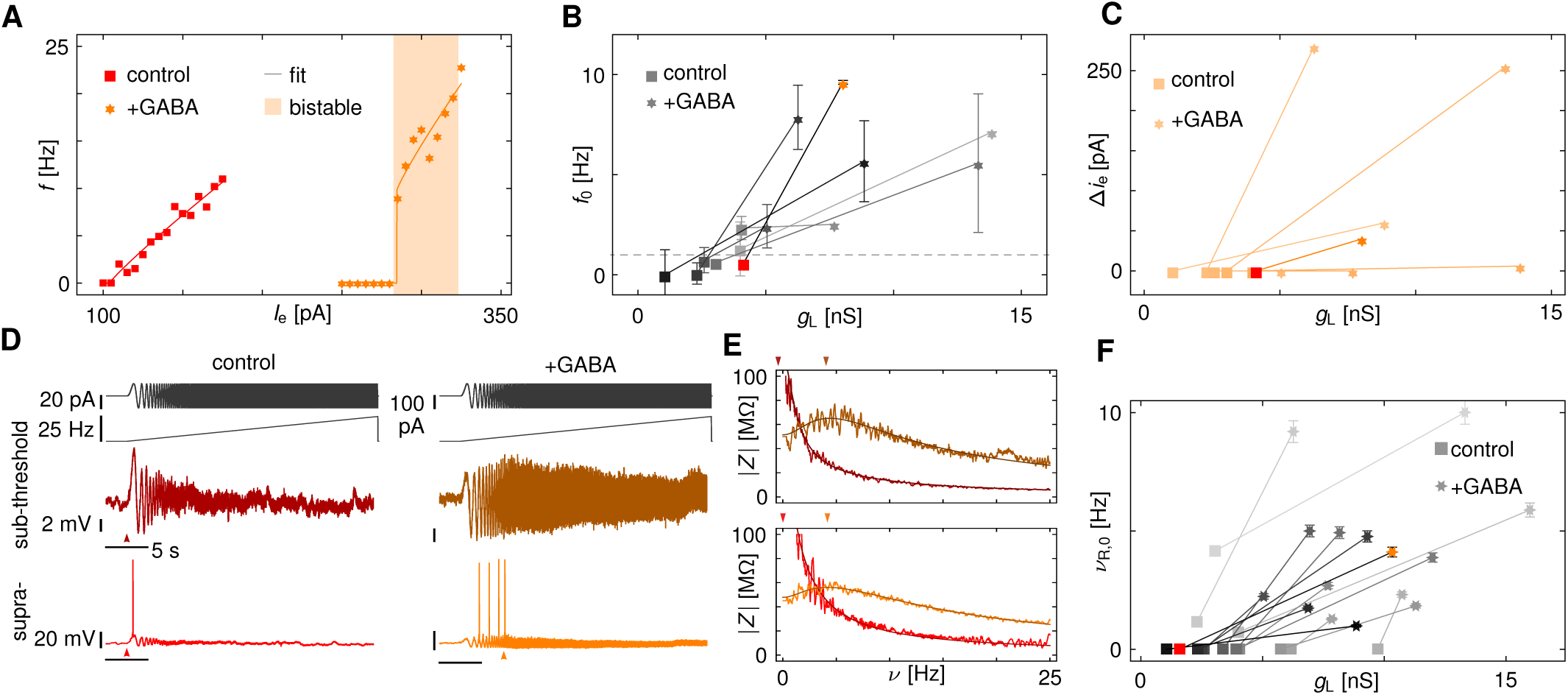
GABA-induced transition from integration to resonance in CA3 pyramidal neurons. (*A*)*f* -*I*-curve and region of bistability of a class-I CA3 neuron without (red) and with application of 500 *μ*M GABA (orange) which switched the excitability from class-I to II. (*B*) Increase in onset firing frequency *f*_0_ due to GABA puff application across different CA3 neurons (*n* = 7). Data from (A) in color. (*C*) Width Δ*I*_e_ of the region of bistability for all cells in (B) before and after puffing 500 *μ*M GABA onto the neurons. In *n* = 5/6 cells, a region of bistability is created, consistent with the general structure of leak-induced excitability transitions (Figs. 1-3). Data from (A) highlighted. (*D*) Voltage responses of the CA3 pyramidal cell in (A) to a ZAP current whose frequency increases linearly from 0 to 25 Hz (top). Responses just below (middle) and above spiking threshold (bottom) for control conditions (left, red) and after puff application of 500 *μ*M GABA (right, orange). (*E*) Impedance curves |*Z*| (line) for the traces in (D) with fit to an LRC circuit (darker line). Arrows indicate resonance frequencies *ν*_R_ obtained from the fit. (*F*) Resonance frequencies *ν*_R,0_ obtained as the average of the resonance frequencies just below and above threshold as a function of *g*_L_ for all cells before and after GABA application. *ν*_R,0_ of all cells (*n* = 15) increased with increasing leak. All non-resonant cells (*n* = 12) became resonant. Data from (D, E) in color.

### Control of Stimulus Encoding and Network Dynamics

How do synaptically controlled changes in the AP dynamics affect neuronal function? In Fig. 7A the resonance properties of a single model neuron is modulated by activation of slower shunting inhibition (as in Fig. 6) which in turn regulates the neuron’s spiking response to constant and oscillatory stimuli. Controlled induction and tuning of resonances thus enables the flexible gating of oscillatory inputs. For CA3 pyramidal neurons the induced resonances have a frequency range of 2 *-* 12 Hz (Figs. 5, 6) and thus may regulate the cell’s participation in 5-12 Hz theta-frequency rhythms in awake animals.

**Fig. 7:**
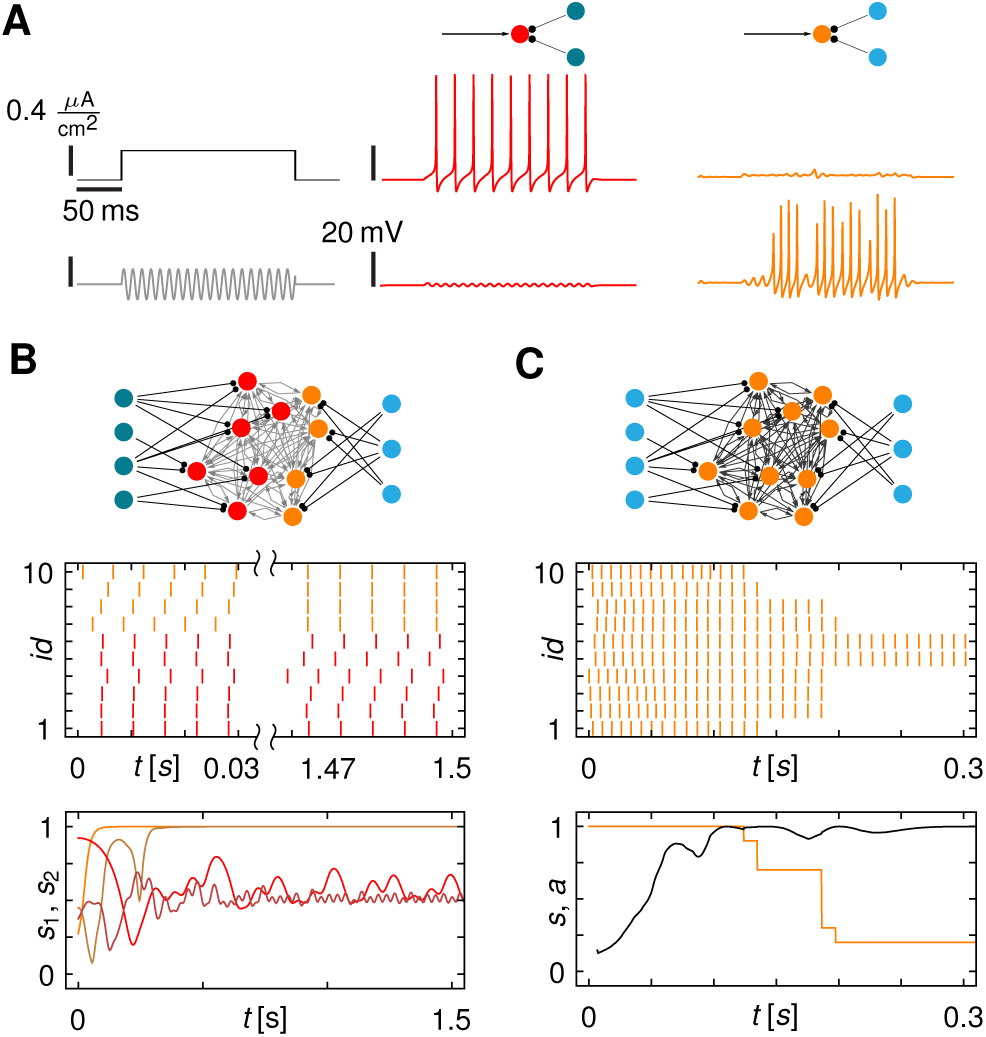
Dynamic control of neuronal excitability and resonance enables variable stimulus encoding and dynamic grouping. (*A*) Class-I excitable model neuron (red or orange disk) receiving external input current (arrow) and synaptic inhibition from inter-neurons (dark or light turquoise disks). For inactive inhibition (middle row) the target neuron is non-resonant and spikes in response to a constant (middle panel) but not to oscillatory input (bottom). Activation of inhibitory synapses (right row) at a Poisson rate of 40 Hz induces resonance in the center neuron (orange disk). Stimulation with an oscillatory stimulus near the resonance frequency induces spiking. Fluctuations in the response are due to stochastic activation of the inhibitory neurons. (*B*) Network (top) of *N* = 10 excitatory neurons (red) receiving synapses from two groups of inhibitory neurons. For silent inhibitory neurons (dark turquoise), the excitatory neurons are class-I (red) and desynchronize (SI Appendix, Fig. S10A). Activating a subset of inhibitory neurons (light turquoise) provides sufficient shunting to dynamically change a subset of postsynaptic neurons to class-II (orange), giving rise to a hybrid class-I/II network. Class-II neurons, initially asynchronous, synchronize (orange) while class-I neurons, initially synchronous, desynchronize (red). *s*_1_ (red line) and *s*_2_ (orange line) are synchrony measures of the individual sub-groups. Random initial conditions lead to the same dynamic grouping (bottom, dark red and orange lines). Switching all neurons to class-II by activation of all inhibitory neurons lead to network wide synchronization (SI Appendix, Fig. S10B). (*C*) Bistability controls the maximal number of synchronized neurons. Network as in (B) with increased coupling weights and all inhibitory neurons active. After an initial synchronization phase, the combined synaptic inputs are sufficiently strong to push neurons from spiking into the basin of attraction of the coexisting fixed point, leaving only a smaller fraction *a* of active neurons (orange line bottom).

Also the collective dynamics of neuronal networks depends on the characteristics of AP generation. For instance, the excitability class of neurons is known to affect their synchronization properties: while class-I neurons tend to desynchronize, class-II neurons synchronize [12]. To see how the different regimes in the BTC structure could be employed to control collective dynamics, we simulated networks of excitatory class-I neurons that were the targets of inter-neurons with slow shunting synapses. For silent inter-neurons, the excitatory neurons are class-I and desynchronize (SI Appendix, Fig. S10A,C). Inhibition effectively increases the leak conductance of the excitatory neurons and switches them to class-II causing them to synchronize (SI Appendix, Fig. S10B,C). Interestingly, if only a subgroup of the excitatory neurons switched to class-II through inhibition, we observe that only the neurons in this sub-ensemble synchronize their spikes. The others become asynchronous (Fig. 7B) despite the stronger synchronous input to all neurons.

More interestingly, positioning neurons into the bistable class-II regime induces synchronization but also limits the total number of active neurons (Fig. 7C): once the synchronized synaptic input pulses become strong enough they push spiking neurons into the basin of attraction of the co-existing resting state (SI Appendix, Fig. S10D). This process continues until the combined pulse strength of the active neurons becomes too weak to silence even more. Together, synchronization and bistability effectively limit the total number of simultaneously active neurons.

Changing the AP dynamics of neurons with shunting inhibition and thereby flexibly shifting their dynamics in the BTC bifurcation structure thus provides a basic biophysical mechanism to control signal processing and collective network dynamics. In particular, this dynamic neuronal excitability can selectively gate oscillatory inputs and bind a self-limited number of neurons by synchrony [32, 33], which has potential roles in flexible neuronal coding and information transmission [34].

### Discussion

Here we systematically investigated the precise structure of transitions in the mechanism of AP generation. We showed that changing both the active, spike generating conductances, as well as the passive leak conductance induces a complex, yet generic, sequence of transitions in regularly firing neurons, from integration to resonance, from uniform dynamics to bi-stable behavior, and from class-I to class-II neuronal excitability (Figs. 1-3, SI Appendix, Fig. S1-S6, Table S1). By tuning the neuron’s capacitance *c*_m_, the leak conductance *g*_L_, and the input current *I*_e_ any class-I neuron can be tuned to a Bogdonav-Takens-Cusp (BTC) bifurcation, for which all sub-transitions collapse into a single point in parameter space. Moving away from this point yields the observed sequence of dynamical transitions. The BTC point underlies and organizes the changes in AP generation and explains the prevalence and complex structure of the transition sequence. We experimentally confirmed the predictions of the BTC-transition structure using dynamic patch-clamp recordings of hippocampal and brain stem neurons. Moreover, activation of GABAergic inhibition moved class-I neurons into different dynamical regimes, making them resonant, bistable, and class-II. Presynaptic activity, therefore, permits dynamical control of single-neuron properties. We linked the regimes in the BTC-transition diagram to different neuronal encoding and filtering properties and collective behaviors in recurrent networks, which shows that not only neuronal dynamics, but also information processing can be flexibly controlled.

The unfolding of the BTC point in the framework of multiple bifurcation theory provides a unified explanation for a number of transition phenomena in AP generation and resonance: applying shunting increases the non-zero onset AP frequency of fast spiking inter-neurons in rat somatosensory cortex neurons [35], and while pyramidal neurons are class-I *in vitro*, they are class-II *in vivo* [20, 36]. Reexamining recordings of neurons in rat somatosensory cortex [35], of pyramidal-and inter-neurons in prefrontal cortex [37] as well as stellate cells of the medial entorhinal cortex [18] reveals signatures of leak-induced bistability. During neuronal development leak changes where shown to adjust AP latency and precision [38].

Resonances and oscillations caused by currents that activate in the sub-threshold regime, such as persistent sodium or hyperpolarization-activated currents, are attenuated in strength and frequency by leak [14, 29, 31]. In contrast, the BTC-structure reveals that leak has the opposite effect on the resonances linked to AP-generation: adding a faster outward current effectively shortens the time scale of rectification until it becomes comparable to the time scale for activating the inward currents and damped oscillations result; the greater the outward conductance, the higher the resonance frequency. Interestingly, the resonances can occur over a wide range of membrane-potentials and can even remain purely sub-threshold, ceasing as the AP threshold is approached. Experimentally, resonances near threshold caused by AP generating currents have been observed [18, 39, 40, 41, 42, 43] as well as increased resonance frequencies due to an increase in outward potassium conductance in phasic vestibular neurons [44].

Dendrites, by effectively imposing an electrical load onto the soma [45], can also affect the excitability class and resonance properties of neurons. Tellingly, for stellate cells in the medial entorhinal cortex, the number of primary dendrites correlates with the resonance frequency [46], and in Purkinje cells switches from class-II to class-I excitability can be observed when pinching the dendritic tree [45].

In contrast to our approach using multiple bifurcation theory, the difference between the two neuronal excitability classes can be explained qualitatively, using the steady-state *I*-*V*-curve [2, 24]:

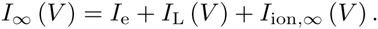

Here *I*_ion,*∞*_ (*V*) are the active ion currents at steady state. Class-I neurons usually have a kink in the steady-state *I*-*V*-curve. The lack of a kink in class-II neurons implies that positive feedback must race against negative feedback to produce an AP; as a consequence, the AP frequency must be nonzero. An increase in the leak conductance flattens out the kink in the steady-state *I*-*V*-curve, and switches the excitability from class-I to II. Thus, heuristically, the critical leak conductance *g*_L,0_ for which the kink disappears is

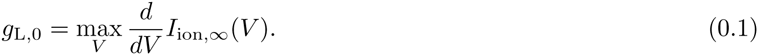

This condition, while sufficient for class-II excitability, is not necessary, however. The positive feedback from a minor inflection in the *I*_*∞*_ (*V*) curve may be too weak to generate slow AP frequencies. Nonetheless, Eq. (0.1) is one of our conditions for the system to be at the BTC point (SI Appendix, Section 2), at which all transitions are localized to a single point in parameter space. Away from the BTC bifurcation, the dynamical transitions occur sequentially at three separate points in the (*I*_e_,*g*_L_)-plane. Neuronal resonance, the switch in excitability, and bistability all appear before an increase in *g*_L_ starts to destabilize damped oscillations at the BT point.

We showed experimentally that activating GABA receptors is sufficient to control excitability and resonance (Fig. 6). Intriguingly, GABA receptors are often preferentially located near the soma [47], which may make them highly effective in dynamically modulating excitability. On longer time scales regulation of potassium leak channels [48] could move neurons into different dynamical regimes of the BTC-structure. In hippocampal pyramidal cells, we found GABA-induced transitions to resonance in the theta band (2 *-* 12 Hz), which could flexibly couple these cells to the oscillations prevalent in this brain area [49]. Up-regulated leak conductances in epileptic brain tissue [50] have the potential to induce a switch from class-I to class-II excitability and thus could further support pathological oscillations and synchronization.

Presynaptic control of postsynaptic neuronal excitability provides a flexible mechanism to regulate responses to oscillatory stimuli and to dynamically group sub-ensembles of neurons and their maximal participation rate in synchronous activity (Fig. 7). Dynamic grouping in turn has potential roles for neuronal coding and information transmission, such as binding by synchrony [33, 51] or communication through coherence [34] and can facilitate top-down processing across network layers [10, 52], underscoring the importance of conductance and excitability for computation in the brain.

## Materials and Methods

### Slice preparation

Brain slices were prepared from Mongolian gerbils (Meriones uniguiculatus) of postnatal day (P) 10 to 18, as described in [53]. In brief, animals were anesthetized, sacrificed, and then brains were removed in cold dissection solution containing (in mM) 50 sucrose, 25 NaCl, 25 NaHCO_3_, 2.5 KCl, 1.25 NaH_2_PO_4_, 3 MgCl_2_, 0.1 CaCl_2_, 25 glucose, ascorbic acid, 3 myo-inositol and 2 Na-pyruvate (pH 7.4 when bubbled with 95% O_2_ and 5% CO_2_). Subsequently, 200 *μ*m thick transverse slices containing the DNLL (P10-11) or 300 *μ*m thick horizontal slices containing the hippocampus (P16-18) were taken with a VT1200S vibratome (Leica). Slices were incubated in extracellular recording solution (same as dissection solution but with 125 mM NaCl, no sucrose, 2 mM CaCl_2_ and 1 mM MgCl_2_) at 36°C for 45 minutes, bubbled with 5% CO_2_ and 95% O_2_.

### Electrophysiology

In the recording chamber, slices were visualized and imaged with a TILL Photonics system attached to a BX50WI (Olympus) microscope equipped with gradient contrast illumination (Luigs and Neumann). All recordings were carried out at near physiological temperature (34 *-* 36*°*C) in current-clamp mode using an EPC10/2 amplifier (HEKA Elektronik). Data were acquired at 50kHz and filtered at 3kHz. The bridge balance was set to 100% after estimation of the access resistance and was monitored repeatedly during recordings. The internal recording solution consisted of (in mM): 145 K-gluconate, 5 KCl, 15 HEPES, 2 Mg-ATP, 2 K-ATP, 0.3 Na_2_-GTP, 7.5 Na_2_- Phospocreatine, 5 K-EGTA (pH 7.2). 100 *μ*M Alexa 488 or 568 were added to the internal solution to control for cell type and location. To apply an artificial extrinsic leak conductance during recordings, an analogue conductance amplifier (SM-1, Cambridge Conductance) applied a constant conductance with a reversal potential equal to the neuron’s resting potential. GABA was puffed at an initial concentration of 500 *μ*M with continuous low pressure controlled by a picospritzer (Picospritzer III, Science Products). Glycinergic and glutamatergic synaptic inputs were blocked with 0.5 *μ*M Strychnine, 20 *μ*M DNQX, 10 *μ*M R-CPP, and GABAergic inputs were blocked with 10 *μ*M SR95531 in all experiments except when GABA was used as an agonist.

### Data Analysis

The leak was estimated from small negative step currents of amplitude *δI*_e_ and 0.5 s duration introduced into the neuron. The average of 50 such traces was fitted by *V* = *V*_min_ + *V*_0_ exp (*-t/τ*_L_). The leak was estimated as *g*_L_ = *δI*_e_*/* (*V*_min_ *- V*_0_), and the capacitance as *c*_m_ = *τ*_L_*g*_L_. To validate the dynamic-clamp method, we determined the relationship between the imposed leak *g*_L,e_ and the measured leak *g*_L_. This relationship was linear with a slope close to 1. The capacitance remained constant for different externally applied leak conductances. One-second long depolarizing step currents were used to determine the f-I-curve and the area of bistability. The AP current threshold was estimated by a semi-automated search using 0.5 s long step currents. Voltage deflections crossing *-*20 mV were classified as APs [54]. For the firing to be classified as periodic, APs had to occur throughout the 1s long stimulation at intervals with a coefficient of variation (CV) less than 50%. Spike frequency f was determined by the average of the inverse inter-spike intervals. If there were fewer than four APs, the inverse latency to the first AP was included in the average. A transition between bistable and periodic APs was deemed to occur when the CV of the inter-spike intervals changed by more than a factor of 1.5 between successive step currents. Clear alternations between periodic and non-periodic APs were seen in the voltage traces in the regimes of bistability. The spike onset frequency *f*_0_ was determined by fitting the *f − I*-curve to *f* (*I*_e_) = Θ (*I*_e_ − *I*_0_) [*α* (*I*_e_ − *I*_0_)^*β*^ + *f*_0_] with fit parameters *f*_0_ *≥* 0, *I*_0_, *α ≥* 0, *β* and Θ a step-like threshold. Standard errors for the parameter estimates were used. Floquet multiplier *λ* were estimated by fitting an exponential decay towards the steady state AP amplitude. Resonance frequencies were estimated by inducing a ZAP stimulus *I*_ZAP_ of the form

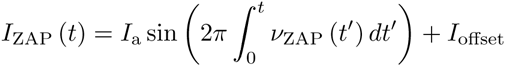

where the time dependent instantaneous frequency *ν*_ZAP_ (*t*) was ramped linearly from 0 to 25 Hz in 30 s and, for controls, from 25 Hz down to 0 Hz in 30 s. The ZAP current amplitude *I*_*a*_ was calibrated before each sweep to yield a voltage deflection of *±*5 mV in the low frequency limit. The constant offset current *I*_offset_ was adjusted to make the cell be either just below or above spike threshold. The frequency-resolved impedance *Z* (*ν*) was estimated from the discrete Fourier transforms of *δI* (*t*) and *δV* (*t*), with 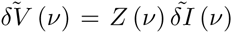. To estimate the resonance frequency, we fitted |*Z* (*ν*) |^*2*^ to the impedance of an RCL circuit |*Z*_RCL_ (*ν*)|^2^= (*a* + *bν*^2^) */ ν*^4^ + *cν*^2^ + *d*, with fit parameters (*a*,*b*,*c, d*). Frequency components of 0.5 Hz and less were dropped to exclude effects of slow drifts [42]. The resonance frequency *ν*_R_ was then determined by arg max *Z*_RCL_ (*ν*).

### Bifurcation Analysis

Bifurcation diagrams and Floquet multipliers [55] were numerically computed using the continuation software AUTO [56]. To simulate model dynamics, we wrote a dynamical systems package for Wolfram’s Mathematica 9 with an interface to AUTO. For the bifurcation diagrams in the main text we used the Wang-Buzsaki neuron model [57]. The same results are obtained for many neuron models including the those by Rinzel [30], Morris-Lecar [58], Erisir et al. [59], Rose-Hindmarsh [60], Connor-Stevens [61], Golomb [62] and reduced pyramidal neuron models [36, 63, 64] (see SI Appendix, Figs. S3-S6).

### Organization of Neuronal Excitability and Resonance Transitions

The impact of leak currents on neuronal excitability was studied using conductance based neuron models of the form

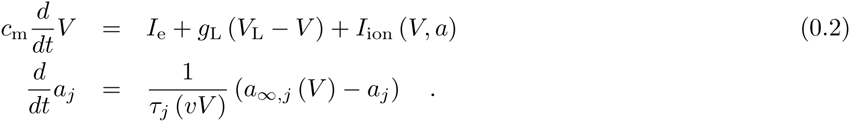

The active ion currents are given by

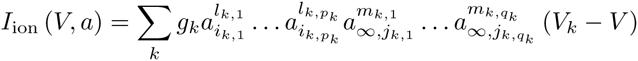

which depend on the maximal conductances *g*_*k*_, reversal potentials *V*_*k*_, and activation variables *a* = (*a*_1_, *a*_2_, *…, a_N_*), steady state activations *a*_*∞*,*j*_ and time constants *τ*_*j*_. Equation (0.2) covers a large number of neuron models, including those in which some activation variables have been replaced by their steady state value *a*_*∞*,*j*_ (*V*). By using a combination of a center-manifold and normal form reduction (62) together with multiple bifurcation theory we prove the following results (see SI Appendix, Section 4 for full proofs):

**Theorem.** Every class-I neuron of the form (0.2) has a Bogdanov-Takens-Cusp point in the parameter space of input, leak conductance, and capacitance.

The proof makes use of two observations: first, the bifurcation parameters *I*_e_, *g*_L_, and *c*_m_ are general parameters occurring in all conductance-based neuron models and only appear as coefficients of terms constant or linear in *V*. Second, the dynamics of the gating variables are coupled solely through the membrane potential *V*.

**Proposition.** If

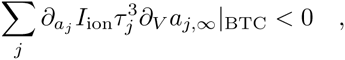

the BTC point is of focus or elliptic type (cf. Figs. 1, 2,S1,S2).

In appropriate coordinates *u*_1_,*u*_2_ the unfolding of the BTC point takes the form

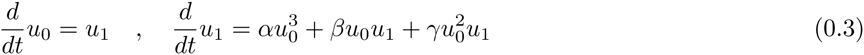

that represents the simplest neuron model that in dependence of parameter *α*,*β* and *γ* captures all the dynamical regimes observed in neuronal excitability transitions. For a neuron model satisfying the conditions of the above theorem a parameter dependent coordinate transformation exists [25] that up to higher order terms transforms it into the equations (0.3) (cf. SI Appendix, Section 4 and Eq. (S.20),(S.26)).

### Linear Response for Conductance-based Models

Neuronal resonance frequencies were determined from the linear response to sinusoidal input stimuli around a steady state *x*_0_ = (*V*_0_, *a*_2,0_, …, *a*_*N*,0_)^T^ using Eq. (0.2). The impedance is

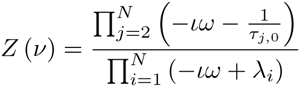

where 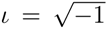 and *λ*_*i*_ are the eigenvalues of the Jacobian *Df* (*x*_0_) (see SI Appendix for details). The transition to resonance was detected by numerical continuation of the condition 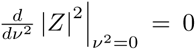. Saddle-point to focus transitions were detected by continuation of zeros of the discriminant for the characteristic polynomial det (*Df* (*x*_0_) *- λ*).

### Network and Resonance Simulations

Networks of *N* Wang-Buzsáki class-I model neurons [57] with *I*_e_ = 4.6 *μ*A*/*cm^2^, *g*_L_ = 0.5 mS*/*cm^2^ and *V*_L_ = *-*60 mV were coupled all-to-all via excitatory synapses. A synapse from pre-synaptic neuron *j* added a current

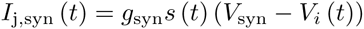

to post-synaptic neuron *i* with membrane potential *V*_*i*_. For weak (strong) excitatory synapses we used *g*_L_ = 0.0025 mS*/*cm^2^ (0.005 mS*/*cm^2^), *V*_syn_ = 0 mV. The activation *s* (*t*) evolved as

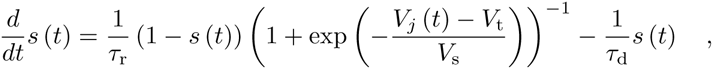

with *τ*_r_ = 0 ms, *τ*_d_ = 1.5 ms, *V*_t_ = *-*20 mV, *V*_s_ = 1 mV. Activation of inhibitory neurons was simulated by Poisson spike trains activating shunting synapses with *τ*_d_ = 5 ms and *V*_syn_ = *-*51.4 mV, resulting in a weakly fluctuating shunting conductance around *g*_*syn*_ = 0.7 mS*/*cm^2^, sufficient to shift neurons to class-II excitability (cf. Figs. 1, 2). Synchrony was measured by the vector strength *s* = |∑ _*j*_ *e*^*ιφi*^| with *φ*_*i*_ the instantaneous phase of neuron *i* estimated by linear interpolation from 0 to 2*π* between spikes. Spikes were determined by positive voltage crossings at *V*_t_ = *-*20 mV.

## Acknowledgements

We thank J. Hudspeth, M. Gutnick and A. Neef for useful discussions. This work has been supported by the German Federal Ministry for Education and Research through the Bernstein Center for Computational Neuroscience Munich (Grant 01GQ0440) and a Fellowship from the Rockefeller University (to C.K.).

## References

[1] Hodgkin A (1948) The local electric changes associated with repetitive action in a non-medullated axon. J Physiol 107(2):165–181.

[2] Rinzel J, Ermentrout G (1989) Analysis of Excitability and Oscillations. Methods in Neuronal Modeling, eds Koch C, Segev I (MIT Press), pp 251–291.

[3] Izhikevich EM (2010) Dynamical Systems in Neuroscience (MIT Press).

[4] St-Hilaire M, Longtin A (2004) Comparison of coding capabilities of Type I and Type II neurons. J Comput Neurosci 16(3):299–313.

[5] Gutkin BS, Ermentrout GB (1998) Dynamics of membrane excitability determine interspike interval variability: a link between spike generation mechanisms and cortical spike train statistics. Neural Comput 10(5):1047–1065.

[6] Gutkin BS, Ermentrout GB, Reyes AD (2005) Phase-response curves give the responses of neurons to transient inputs. J Neurophysiol 94(2):1623–1635.

[7] Tateno T, Robinson H (2006) Rate coding and spike-time variability in cortical neurons with two types of threshold dynamics. J Neurophysiol 95(4) 2650–2663.

[8] Broicher T, et al. (2012) Spike phase locking in CA1 pyramidal neurons depends on background conductance and firing rate. J Neurosci 32(41):14374–14388.

[9] Schleimer JH, Stemmler M (2009) Coding of Information in Limit Cycle Oscillators. Phys Rev Lett 103(24):248105.

[10] Lengyel M, Kwag J, Paulsen O, Dayan P (2005) Matching storage and recall: hippocampal spike timingdependent plasticity and phase response curves. Nat Neurosci 8(12), 1677–1683.

[11] Hansel D, Mato G, Meunier C (1995) Synchrony in excitatory neural networks. Neural Comput 7(2):307–237.

[12] Ermentrout B (1996) Type I membranes, phase resetting curves, and synchrony. Neural Comput 8(5):979–1001.

[13] Ratté S, Hong S, De Schutter E, Prescott SA (2013). Impact of neuronal properties on network coding: roles of spike initiation dynamics and robust synchrony transfer. Neuron 78(5):758–772.

[14] Hutcheon B, Yarom Y (2000) Resonance, oscillation and the intrinsic frequency preferences of neurons. Trends Neurosci 23(5):216–222.

[15] Llinás R (1988) The intrinsic electrophysiological properties of mammalian neurons: insights into central nervous system function. Science 242(4886):1654–1664.

[16] Buzsáki G, Draguhn A (2004) Neuronal oscillations in cortical networks. Science 304(5679):1926–1929.

[17] Vaidya SP, Johnston D (2013) Temporal synchrony and gamma-to-theta power conversion in the dendrites of CA1 pyramidal neurons. Nat Neurosci 16(12):1812–20.

[18] Alonso A, Klink R (1993) Differential electroresponsiveness of stellate and pyramidal-like cells of medial entorhinal cortex layer II. J Neurophysiol 70(1):128–143.

[19] Haas JS, White JA (2002) Frequency selectivity of layer II stellate cells in the medial entorhinal cortex. J Neurophysiol 88(5):2422–2429.

[20] Prescott SA, Ratté S, De Koninck Y, Sejnowski TJ (2006) Nonlinear interaction between shunting and adaptation controls a switch between integration and coincidence detection in pyramidal neurons. J Neurosci 26(36):9084–9097.

[21] Stiefel KM, Gutkin BS, Sejnowski TJ (2008) Cholinergic neuromodulation changes phase response curve shape and type in cortical pyramidal neurons. PLoS One 3(12):e3947.

[22] Yang J, et al. (2009) Membrane current-based mechanisms for excitability transitions in neurons of the rat mesencephalic trigeminal nuclei. Neuroscience 163(3):799–810.

[23] Liu Y, Yang J, Hu S (2008) Transition between two excitabilities in mesencephalic V neurons. J Comput Neurosci 24(1):95–104.

[24] Prescott SA, De Koninck Y, Sejnowski TJ (2008) Biophysical basis for three distinct dynamical mechanisms of action potential initiation. PLoS Comput Biol 4(10):e1000198.

[25] Dumortier F, Roussarie R, Sotomayor J, Zoladek H (1991) Bifurcations of Planar Vector Fields: Nilpotent Singularities and Abelian Integrals (Springer, New York).

[26] Kuznetsov YA (2005) Practical Computation of Normal Forms on Center Manifolds At Degenerate Bogdanov-Takens Bifurcations. Int J Bif Chaos 15(11):3535–3546.

[27] Guckenheimer J (1986) Multiple bifurcation problems for chemical reactors. Physica D 20(1):1–20.

[28] Guckenheimer J, Holmes P (1983) Nonlinear Oscillations, Dynamical Systems, and Bifurcations of Vector Fields (Springer, New York).

[29] Fernandez FR, White JA (2008) Artificial synaptic conductances reduce subthreshold oscillations and periodic firing in stellate cells of the entorhinal cortex. J Neurosci 28(14):3790–3803.

[30] Rinzel J (1985) Excitation dynamics: insights from simplified membrane models. Fed Proc 44(15):653–675.

[31] Hutcheon B, Miura RM, Puil E (1996) Models of subthreshold membrane resonance in neocortical neurons. J Neurophysiol 76(2):698–714.

[32] Hebb DO (1949) The organization of behaviour (John Wiley & Sons).

[33] von der Malsburg C. The correlation theory of brain function. Intern. Rep. 81–2, Max-Planck-Institute for Biophysical Chemistry, Göttingen, Germany.

[34] Fries P (2005) A mechanism for cognitive dynamics: neuronal communication through neuronal coherence. Trends Cogn Sci 9(1):474–480.

[35] Tateno T, Harsch A, Robinson HPC (2004) Threshold firing frequency-current relationships of neurons in rat somatosensory cortex: type 1 and type 2 dynamics. J Neurophysiol 92(4):2283–2294.

[36] Prescott SA, Ratté S, De Koninck Y, Sejnowski TJ (2008) Pyramidal neurons switch from integrators in vitro to resonators under in vivo-like conditions. J Neurophysiol 100(6):3030–3042.

[37] Fellous JM, et al. (2001) Frequency dependence of spike timing reliability in cortical pyramidal cells and interneurons. J Neurophysiol 85(4):1782–1787.

[38] Franzen DL, et al. (2014) Development and modulation of intrinsic membrane properties control the temporal precision of auditory brainstem neurons. J Neurophysiol 113(3):524–536.

[39] Alonso A, Llinas RR (1989) Subthreshold Na+-dependent theta-like rhythmicity in stellate cells of entorhinal cortex layer II. Nature 342(6246):175–177.

[40] Gutfreund Y, Yarom Y, Segev I (1995) Subthreshold oscillations and resonant frequency in guinea-pig cortical neurons: physiology and modelling. J Physiol 483(3):621–640.

[41] Hu H, Vervaeke K, Storm JF (2002) Two forms of electrical resonance at theta frequencies, generated by Mcurrent, h-current and persistent Na+ current in rat hippocampal pyramidal cells. J Physiol 545(1):783–805.

[42] Erchova I, Kreck G, Heinemann U, Herz AVM (2004). Dynamics of rat entorhinal cortex layer II and III cells : characteristics of membrane potential resonance at rest predict oscillation properties near threshold. J Physiol 560(1):89–110.

[43] Prescott SA, De Koninck Y (2009) Impact of background synaptic activity on neuronal response properties revealed by stepwise replication of in vivo-like conditions in vitro. Dynamic-Clamp, eds Bal T, Destexhe A (Springer), pp 89–114.

[44] Beraneck M, et al. (2007) Differential intrinsic response dynamics determine synaptic signal processing in frog vestibular neurons. J Neurosci 27(16):4283–4296.

[45] Bekkers JM, Häusser M (2007) Targeted dendrotomy reveals active and passive contributions of the dendritic tree to synaptic integration and neuronal output. Proc Natl Acad Sci USA 104(27):11447–11452.

[46] Garden DLF, et al. (2008) Tuning of synaptic integration in the medial entorhinal cortex to the organization of grid cell firing fields. Neuron 60(5):875–889.

[47] Buhl E, Halasy K, Somogyi P (1994) Diverse sources of hippocampal unitary inhibitory postsynaptic potentials and the number of synaptic release sites. Nature 368(6474):823–828.

[48] Brickley SG, Revilla V, Cull-Candy SG, Wisden W, Farrant M (2001) Adaptive regulation of neuronal excitability by a voltage-independent potassium conductance. Nature 409(6816):88–92.

[49] Harris K, Henze D, Hirase H (2002) Spike train dynamics predicts theta-related phase precession in hippocampal pyramidal cells. Nature 417(6890):2116–2118.

[50] Stegen M, Young CC, Haas CA, Zentner J, Wolfart J. Increased leak conductance in dentate gyrus granule cells of temporal lobe epilepsy patients with Ammon’s horn sclerosis. Epilepsia 50(4):646–53.

[51] Singer W, Gray C (1995) Visual feature integration and the temporal correlation hypothesis. Annu Rev Neurosci 18:555–586.

[52] Engel AK, Fries P, Singer W (2001) Dynamic predictions: oscillations and synchrony in top-down processing. Nat Rev Neurosci 2(10):704–716.

[53] Ammer JJ, Grothe B, Felmy F. Late postnatal development of intrinsic and synaptic properties promotes fast and precise signaling in the dorsal nucleus of the lateral lemniscus. J Neurophysiol 107(4):1172–1185.

[54] Todd BS, Andrews DC (1999) The identification of peaks in physiological signals. Comput Biomed Res 32(4):322–335.

[55] Cesari L (1971) Asymptopic behaviour and stability problems in ordinary differential equations. (Springer, New York).

[56] Doedel EJ, et al. (2007) Auto-07p: Continuation and bifurcation software for ordinary differential equations, http://indy.cs.concordia.ca/auto.

[57] Wang XJ, Buzsáki G (1996) Gamma oscillation by synaptic inhibition in a hippocampal interneuronal network model. J Neurosci 16(20):6402–6413.

[58] Morris C, Lecar H (1981) Voltage oscillations in the barnacle giant muscle fiber. Biophys J 35(1):193–213.

[59] Erisir A, Lau D, Rudy B, Leonard CS (1999) Function of specific K+ channels in sustained high-frequency firing of fast-spiking neocortical interneurons. J Neurophysiol 82(5):2476–2489.

[60] Rose R, Hindmarsh J (1989) The assembly of ionic currents in a thalamic neuron I. The three-dimensional model. Proc R Soc London Ser B 237(1288):267–288.

[61] Connor J, Stevens C (1971) Voltage clamp studies of a transient outward membrane current in gastropod neural somata. J Physiol 213(1):21–30.

[62] Golomb D, et al. (2007) Mechanisms of firing patterns in fast-spiking cortical interneurons. PLoS Comput Biol 3(8):e156.

[63] Traub RD, Miles R (1991) Neuronal Networks of the Hippocampus (Cambridge University Press).

[64] Ermentrout GB, Kopell N (1998) Fine structure of neural spiking and synchronization in the presence of conduction delays. Proc Natl Acad Sci USA 95(3):1259–1264.

